# Phylogenetic, geographic and habitat distribution of the green-brown polymorphism in European orthopterans

**DOI:** 10.1101/2020.03.31.016915

**Authors:** Holger Schielzeth

**Author notes:** Address for correspondence: Holger Schielzeth, Population Ecology Group, Institute of Ecology and Evolution, Friedrich Schiller University Jena, Dornburger Straße 159, 07743 Jena, Germany. The authors wish to be identified to the reviewers. Data availability: Data will be made available upon publication of the manuscript. Code and data availability: https://github.com/hschielzeth/OrthopteraPolymorphism.

## Abstract

The green-brown polymorphism among polyneopteran insects represents one of the most penetrant color polymorphisms in any group of organisms. Yet systematic overviews are lacking. I here present analyses of the phylogenetic, geographic and habitat distribution of the green-brown polymorphism across the complete European orthopteran fauna. Overall, 30% of European orthopterans are green-brown polymorphic. Polymorphic species are scattered across the entire phylogenetic tree, including roughly equal proportions of Ensifera and Caelifera. A few taxonomic groups, however, include only brown species. Polymorphic species occur more frequently in clades that contain monomorphic green species than in those without green species. The relative abundance of color morphs in polymorphic species is skewed towards green, and in particular rare/exceptional brown morphs are more common in predominantly green species than rare/exceptional green morphs in predominantly brown species. The patterns of abundances support the hypothesis that loss-of-function mutations play a role in creating polymorphic populations from green species. Polymorphic species are particularly common in moist to mesic grasslands, alpine and arboreal habitats. Dry, open, rocky and cave habitats as well as nocturnal lifestyles are dominated by monomorphic brown species. The proportion of polymorphic species increases from southern to northern latitudes. These marked habitat-dependencies also show that coloration is affected by natural selection and/or environmental filtering. Overall, the results illustrate that the occurrence of the polymorphism is phylogenetically, geographically and ecologically widespread and they suggest that polymorphism is thus potentially in mutation-selection balance across a large number of species.

## Introduction

Intraspecific color polymorphisms have fascinated biologists for a long time. Among the most significant findings, they have spurred the discovery of Mendelian inheritance (Mendel 1866), have stimulated models on polygenic inheritance (Fisher 1930), shifting balance theory and isolation-by distance (Wright 1945), have demonstrated rapid adaptation to environmental change (Kettlewell 1958) and the discovery of a shared genetic basis of phenotypic variation across diverse taxa (Hoekstra 2006). Color polymorphisms represent a particularly conspicuous case of intra-specific diversity and the question is when they arise and how they are maintained. Color polymorphisms are very widespread across a large number of taxa, and each case may have special conditions that create and maintain the polymorphism (Majerus 1998). In some cases, color polymorphisms arise from admixture of divergent populations or shifts from one monomorphic state to another (“transient polymorphisms”, Ford 1966). Such polymorphisms represent snapshots in time and may be transient, without the need to invoke any special mechanisms actively favoring the coexistence of discrete color variants. In other cases, color polymorphisms seem to be actively maintained by balancing selection (“balanced polymorphisms”, Ford 1966).

Widespread color polymorphisms include melanism in animals (Majerus 1998) and corolla color polymorphisms in flowering plants (Rausher 2008). Melanism occurs across a wide range of species such as among birds, mammals, lizards, lepidopterans and ladybirds. However, the cases are rather spread out and not very penetrant within any large group of organisms. Among birds, for example, only about 3.5% of the species are polymorphic for melanistic variants with highest prevalence of about 33% among owls and nightjars (Galeotti et al. 2003; Roulin 2004). Similarly, about 10% of British macro-lepidopterans show melanistic polymorphisms (Kettlewell 1956). Melanism in lepidopterans and ladybirds comes in many forms and many represent geographical clines rather than the local stable coexistence of discrete variants (Majerus 1998). Corolla polymorphisms, mostly involving colored and white variants, have long been discussed (Darwin 1876), although I am not aware of any systematic review that provides specific estimates on how widespread the polymorphism is (but see Ackerman et al. 2011). Here I focus on the green-brown color polymorphism in orthopterans that, I think, outcompetes melanism and corolla coloration for one of most penetrant color polymorphisms in any groups of organisms.

The green-brown polymorphism of grasshoppers represents a long known, but understudied, case of a widespread color polymorphism (Dearn 1990; Rowell 1972). The co-occurrence of green and brown morphs within local populations of single species occurs in both major suborders of Orthoptera, Ensifera and Caelifera, that have separated about 355 Mya (Song et al. 2020). Moreover, the color polymorphism is not limited to orthopterans, but is shared more widely among polyneopteran insects (e.g. phasmids, mantises and gladiators) that shared a common ancestor as long as 415 Mya (Song et al. 2020). Grasshoppers have been model systems for the study of polymorphism, including phase polymorphisms (solitary and gregarious morphs within the same species, Pener and Yerushalmi 1998), pattern polymorphisms (Ahnesjö and Forsman 2003; Nabours 1929) and melanism (Forsman 2011; Peralta-Rincon et al. 2017). However, surprisingly few studies have focused on the green-brown polymorphism. Rowell (1972) reports that about 45% of east African acridid grasshoppers are green-brown polymorphic. In the tropical region the green-brown polymorphism appears more often in seasonal grasslands than in tropical forests and wetlands (Rowell 1972). For temperate regions, it has been found that among British orthopterans green morphs are more abundant in moist habitats as compared to brown morphs dominating in dry habitats (Gill 1981a), illustrating some habitat-dependency of the green-brown polymorphism.

Here I present an analysis of the phylogenetic, geographical and habitat distribution of the green-brown polymorphism across the complete European fauna of orthopterans. The analysis is explorative in the sense of being the first to review the green-brown polymorphism across the entire fauna of a large group of organisms for an entire continent. It is also meant to be hypothesis-generating with respect to habitat and geographical patterns. However, I also test specific hypothesis, in particular the suggestion that polymorphisms favor speciation (West-Eberhard 1986). My main focus, however, is on the question if the green-brown polymorphism arises primarily from in brown species or in green species. Data from species of gomphocerine grasshoppers suggest that the polymorphism is determined by one or few genes with a green allele being dominant over a brown allele (Gill 1981b; Schielzeth and Dieker 2020; Winter et al. 2021), which suggest a role of loss-of-function mutations. I therefore evaluate the relative abundance of green and brown morphs as well as associations of polymorphic species with brown or green species within clades. Skew towards green and preferential co-occurrence of polymorphic with green specie would support this suggestion.

Besides documenting the high prevalence of the green-brown polymorphism, the data show some habitat-dependency and thus environmental filtering or habitat-dependent selection. Skew in the abundance distribution towards green and the phylogenetic clustering of polymorphic with green species are consistent with the hypothesis that the brown morph arises from loss-of-function mutations from a functional green pigmentation pathway. Within populations, the polymorphism might well be maintained by balancing selection, as suggest by its wide distribution across species. The data show evidence that more widespread species are more often polymorphic, but there is no relationship between the green-brown polymorphism and species richness.

## Materials and methods

### Species compilation

I compiled data for all European orthopteran species, excluding Cyprus (but including the Greek islands) and excluding the oceanic islands of Madeira, Canaries and Azores. Species were initially derived from the checklist of European species compiled by Heller et al. (1998). However, the list is rather incomplete in many regions (in particular for the Iberian peninsula) and species were added from the website Grasshoppers of Europe (Adriaens et al. 2019). The geographic scope of Heller et al. (1998) is a little wider in the East, such that a few species from the northern Caucasus are included in Heller et al. (1998), but not Adriaens et al. (2019), and are included here. In all cases, species status was checked with the Orthoptera Species File (Cigliano et al. 2019) and all doubtful species and synonyms were excluded. In total, I compiled an exhaustive list of 1086 species.

### Taxonomy

Taxonomy follows the Orthoptera Species File (Cigliano et al. 2019), including decisions about species status, subgenus, genus, tribe, subfamily, family, superfamily and suborder. In a few cases, some taxonomic levels were missing. For example, there is no official tribe name for the single genus in the subfamily Troglophilinae. A generic tribe was assigned in this case. In most cases, there was no subdivision into subgenera and in most of the remaining cases, all species of a genus were assigned to subgenera. In very few cases, a genus was split into subgenera, with a few species being unassigned (4 species in *Chorthippus*). These were treated as belonging to a separate (unnamed) subgenus.

### Distribution

Distributional information was extracted from Heller et al. (1998), with the modification that I separated Greece from the rest of south-eastern Europe: This resulted in 13 distinct regions (1) British Isles and Island, (2) Norway and Sweden, (3) Finland, Estonia, Latvia and northern Russia (north of 58°N), (4) Netherlands, Belgium, Luxemburg and mainland France, (5) Central Europe, including Denmark, Germany, Poland, Czech Republic, Slovakia, western Ukraine and the northern parts of Switzerland and Austria, (6) Eastern Europe including Lithuania, Belarus and the central parts of European Russia (about 50-58°N), (7) Iberian Peninsula including Balearic Islands, (8) Italy, Corsica, Malta and the southern parts of Switzerland and Austria, (9) Balkan states and SE Europe, including Hungary, Romania, Bulgaria and the European part of Turkey, (10) Greece, (11) Eastern Ukraine and southern Russia south of about 50°N, but excluding the Caucasus region, (12) European parts of Kazakhstan and adjacent parts of Russia, (13) northern Caucasus region. The assignment to regions follows Heller et al. (1998) except for a few obvious corrections, such as in cases of taxonomic reassignments. Missing species (most of which rather localized) were assigned based on distributional information in the Orthoptera Species File (Cigliano et al. 2019). Species from Greece were assigned based on the recent Fauna of Greece (Willemse et al. 2018). Since Heller et al. (1998) combined the entire south-east European region (including Greece), species endemic to Greece were treated as missing from the Balkan region. I used the number of the 13 regions in which a species occurs as a measure of its range size.

### Scoring of color morphs

Color morphs were scored in adult individuals based on images from trusted sources (see below) and from own field observations. Green morphs were identified based on areas of clear green color, while all other individuals were scored as brown (see Figure 1 for examples). Note that brown encompasses a range of non-green colors, including gray, black and reddish. The separation into green versus non-green (brown) was motivated by the hypothesis that there are specific pigments that produce green coloration and that the two groups differ in the presence/absence of these pigments. Categorization was straightforward in most cases. In a few cases, however, categorization was more difficult. There are three main groups of these: (i) Some species of Pholidopterini show a yellowish underside that might sometimes appear as a greenish tinge. In such cases, only individuals with unambiguously green color were scored as green. (ii) In some species (such as some *Barbitistes* and *Poecilimon*) there is a range from clear green phenotypes with variable black markings to almost entirely black individuals. Only individuals in which there was no green areas left were scored as brown. (iii) In some species (such as species of *Arcyptera*) the body surface is very shiny and color sometimes difficult to judge. However, in most of these cases, it was possible to find unambiguously green and unambiguously brown individuals (hence most of these species are polymorphic). Aside from cases (i) and (ii), in which the green-brown polymorphism could be considered gradual and continuous, variation was almost exclusively discrete. Color morph classification was possible for 949 species (87%).

**Figure 1:**
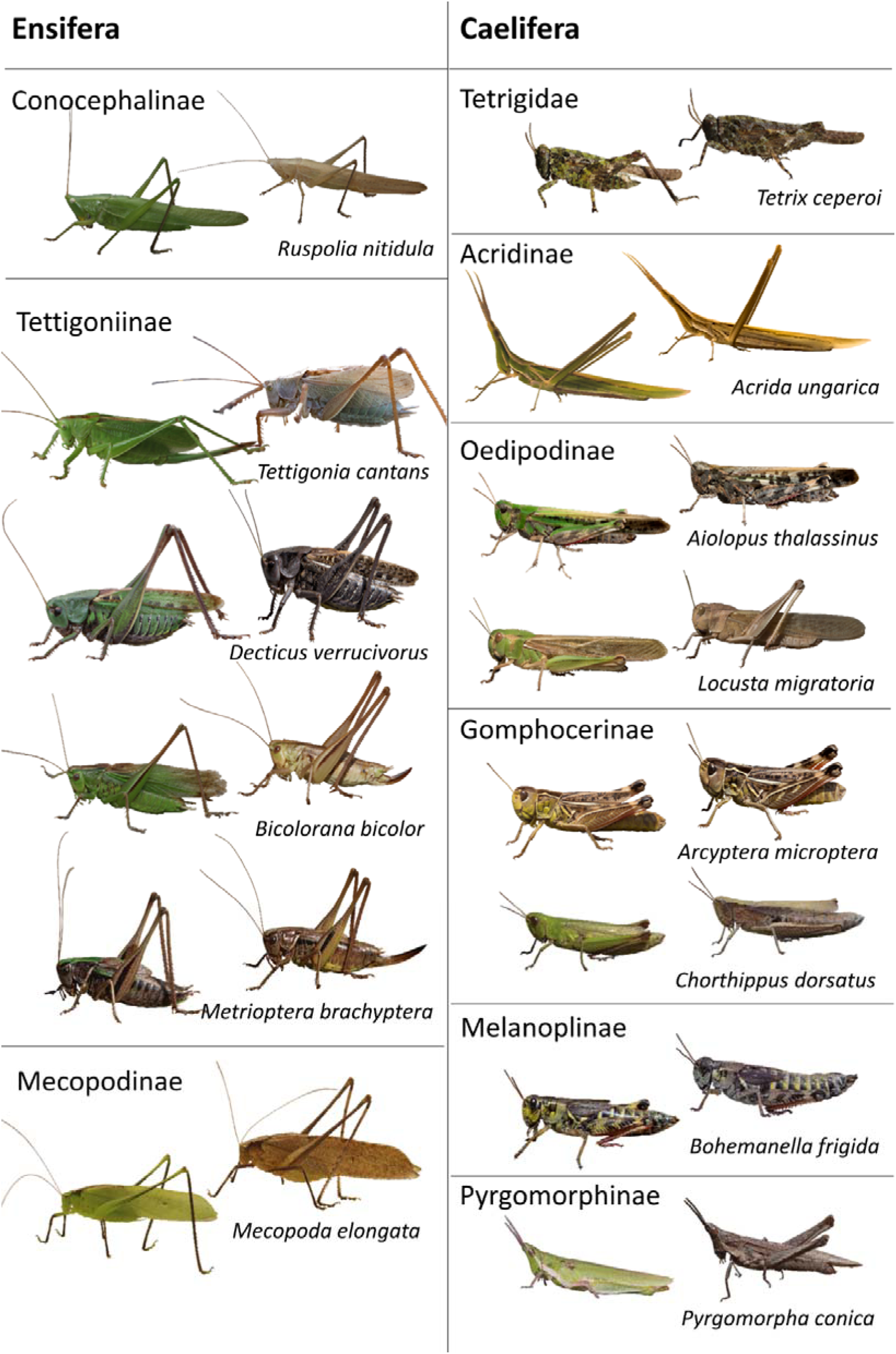
Examples of polymorphic orthopteran species from different subfamilies. The panel shows a mix of sexes, but morph identity is not sex-limited in any of the cases shown. The cases of *Tettigonia cantans* and *Bicoloriana bicolor* illustrate cases where faded brown variants occur at low frequency in green populations. *Metrioptera brachyptera* shows a case where green is limited to the dorsal side. *Tetrix ceperoi* shows a case where green morphs occur at relatively low frequency in brown populations. *Arcyptera microptera* illustrates a case where the body surface is shiny and colors sometimes more difficult to judge. The species *Mecopoda elongata* does not occur in Europe and is added as another example for the green-brown polymorphism in Ensifera.

### Polymorphism survey

Polymorphisms were scored from images (and in a few cases verbal descriptions) in field guides and faunas from Catalonia (Olmo Vidal 2006), France (Bellmann and Luquet 2009; Sardet et al. 2015), Switzerland (Baur et al. 2006), Austria (Zuna-Kratky et al. 2017), Germany (Fischer et al. 2016), Central Europe (Bellmann et al. 2019), Italy (Iorio et al. 2019), Greece (Willemse et al. 2018) and eastern Romania (Iorgu and Iorgu 2008). Furthermore, I searched the websites Grasshoppers of Europe (www.grasshoppersofeurope.de, Adriaens et al. 2019), Pyrgus (http://www.pyrgus.de, Wagner 2019) and Orthoptera.ch (Roesti and Rutschmann 2019) for further images. These websites are run by experts and provide reliable species identifications.

Finally, I searched images on flickr and google based on scientific species names. Flickr provides search results only from the figure caption and these turned out to give very reliable species identification, likely because only people with sufficient expertise would provide scientific species names. Google searches provide fuzzier matching and usually showed some well-identified results followed by a long series of other images. Critical photographs were carefully evaluated by visiting the website, checks of source and location, as well as with my own taxonomic experience on orthopterans. Finally, I added my own field experience, which covers mostly species from central and western Europe. It might seem useful to search museum collections for the green-brown polymorphism. However, the green colors are often lost during preservation (e.g. the green pigments appear to be soluble in ethanol) and are subject to fading, such that I consider scoring from museum collections unreliable.

### Relative morph abundances

One of the main interests of the compilation was in assessing whether there is an association of polymorphic species with either brown or green populations. I therefore compiled the relative abundance of the two morphs for all species. Morph abundances were estimated from field guide information, image searches and own experience and classified into seven categories: monomorphic (100%), dominant (>50%), common (20-50%), regular (10-20%), rare (2-10%) and exceptional (<2%). Being based largely on image searches, these categorizations cannot be exact. However, the broad categorization was relatively easy in most cases, with the following exceptions: (i) in species with two very common morphs it is sometimes difficult to judge which one is more common (assigned dominant) and which one is less common (assigned common). Misclassifications between these two categories are therefore expected. (ii) For species with very rare occurrence of one of the morphs and only a few images available, exceptional or rare morphs could well be overlooked. These were nevertheless categorized as absent, such that the number of polymorphic species is probably underestimated. (iii) The categorization of rare vs. exceptional might seem difficult, but turned out to be relatively easy. The category ‘exceptional’ was used only when long series of images resulted in only single images of the rare morph. This applied to species like *Phaneroptera falcata* (which is typically bright green) or *Oedipoda caerulescens* (which is typically brown, grey or black). It is therefore more likely that species that show exceptional phenotypes are classified as absent rather than being misassigned as rare. It is possible that in a few cases morphs categorized as rare are in fact be rather exceptional.

### Habitat categorization

Habitat preferences were compiled from descriptions in the field guides and faunas mentioned above. I defined the following broad categories: (1) dunes and beaches, (2) pond and river margins (including temporary flooded areas) (3) marshes and reed beds, (4) moist grasslands (including a few tundra species), (5) mesic grasslands, (6) dry grasslands, (7) grasslands with abundant tall herbs, (8) rural areas with herbs, shrubs and open ground, (9) open ground, (10) shrubs and heath, (11) mesic bushland, (12) dry bushland, (13) forest margins, (14) woodlands (including clearings), (15) subalpine and alpine shrubs, (16) subalpine and alpine grasslands, (17) alpine rocky habitats, (18) lowland rocky habitats, (19) stony walls, (20) domestic habitats, and (21) caves. I aimed to assign all species to habitats in which they occur regularly. Habitat information could be compiled for 670 species (62%). I used the number of the 21 habitat classes in which a species occurs as a measure of habitat diversity and ultimately the width of a species’ ecological niche.

### Habitat use

I categorized relevant aspects of habitat use in five categories: (1) *Subterranean* for species living underground, under logs or stones, or in caves, (2) *ground-dwelling* for species typically locating themselves on the ground, (3) *mixed* for species that partly perch on the ground, partly in the vegetation, and (4) *vegetation-dwelling* for species predominately locating themselves and moving in dense herbal vegetation, low shrubs, or grasses, (5) *bushes and trees* for species typically living in higher vegetation layers. Habitat information could be compiled for 680 species (63%).

Furthermore, I classified the activity pattern into three categories: (1) nocturnal for all species almost exclusively active at night, (2) active during day and night, in particular including species that display at night, but are also active during the day and thus exposed to diurnal predators, (3) diurnal species predominately active during the day. Diurnality categories were mutually exclusive and could be compiled for 673 species (62%). Finally, I categorized seasonal activity of imagoes into (1) spring (April-June), (2) summer (late June-November), and (3) winter (species that overwinter as imagoes). Species active in multiple seasons were assigned to multiple categories. Seasonality information could be compiled for 688 species (63%).

### Statistical analyses

It has been suggested that polymorphisms are positively associated with speciation. I therefore regress the number of species within taxonomic groups on the proportion of polymorphic species within these groups. Similarly, I regress the proportion of polymorphic species on the species range size and habitat diversity. Furthermore, I use χ tests to test whether the occurrence of rare morphs is more common in predominantly green or predominately brown species, and I use χ tests for whether polymorphic species occur more frequently in clades with otherwise only monomorphic green, otherwise only monomorphic brown or both types of monomorphic species. All data processing and statistical analysis were done in R 4.0.2 (R Core Team 2020).

One might argue that phylogentic control is necessary in the analysis. However, there is no well-resolved phylogeny for European orthopterans, certainly not at the species-level resolution that is required for the analysis. Any phylogenetic analysis (or ancestral state reconstruction) hinges on accurate phylogenetic information otherwise it will be uninformative to due to known biases. But I also argue that phylogenetic control is not strictly necessary for the patterns that I describe here. Associations with the geographic or habitat distribution might well contain a phylogenetic signal, but association are ecologically relevant in any case. With respect to evolutionary questions, the scattered distribution across the phylogeny clearly illustrated that the green-brown polymorphisms is not an isolated phenomenon that is limited to few clades. Still, a phylogenetic pattern is evident in that some groups are fixed for brown morphs as I show and discuss below. The analysis of associations with green/brown species as conduced at different taxonomic levels (general and tribes) and finding similar patterns in these analyses illustrates that the associations are largely independent of phylogeny. It is thus mostly the analysis of relative abundance that might be affected by phylogeny and that might thus be treated with caution – which is anyway advisable because of the coarse categorization.

## Results

In total, 949 species (87%) could be reliably assigned a color morph classification. Among classified species, 51% are monomorphic brown, 19% monomorphic green, and 30% polymorphic. Excluding the 165 almost exclusively brown crickets (Grylloidea and Gryllotalpoidea, with one polymorphic exception) and cave crickets (Rhaphidophoroidea, with one polymorphic exception), left 42% of monomorphic brown species, 22% monomorphic green species, and 36% polymorphic species.

### Phylogenetic distribution

Both suborders, Ensifera and Caelifera, contain many polymorphic species (29% in Ensifera, 33% in Caelifera, Figure 2). Aside from the (almost) exclusively brown crickets (Grylloidea), mole crickets (Gryllotalpoidea), cave crickets (Rhaphidophoroidea), and pygmy mole crickets (Tridactyloidea), all superfamilies include polymorphic species (Figure 2). All subfamilies of the single family Tettigoniidae of Tettigonioidea have polymorphic species (Figure 2) as well as all families of Acridoidea (Figure 2). Only at the level of subfamilies there is a divide with a few smaller subfamilies (besides crickets) being exclusively brown (Calliptaminae [9 species], Pezotettiginae [3 species], Thrinchinae [11 species], Dericorythinae [2 species], Dentridactylinae [1 species]).

**Figure 2:**
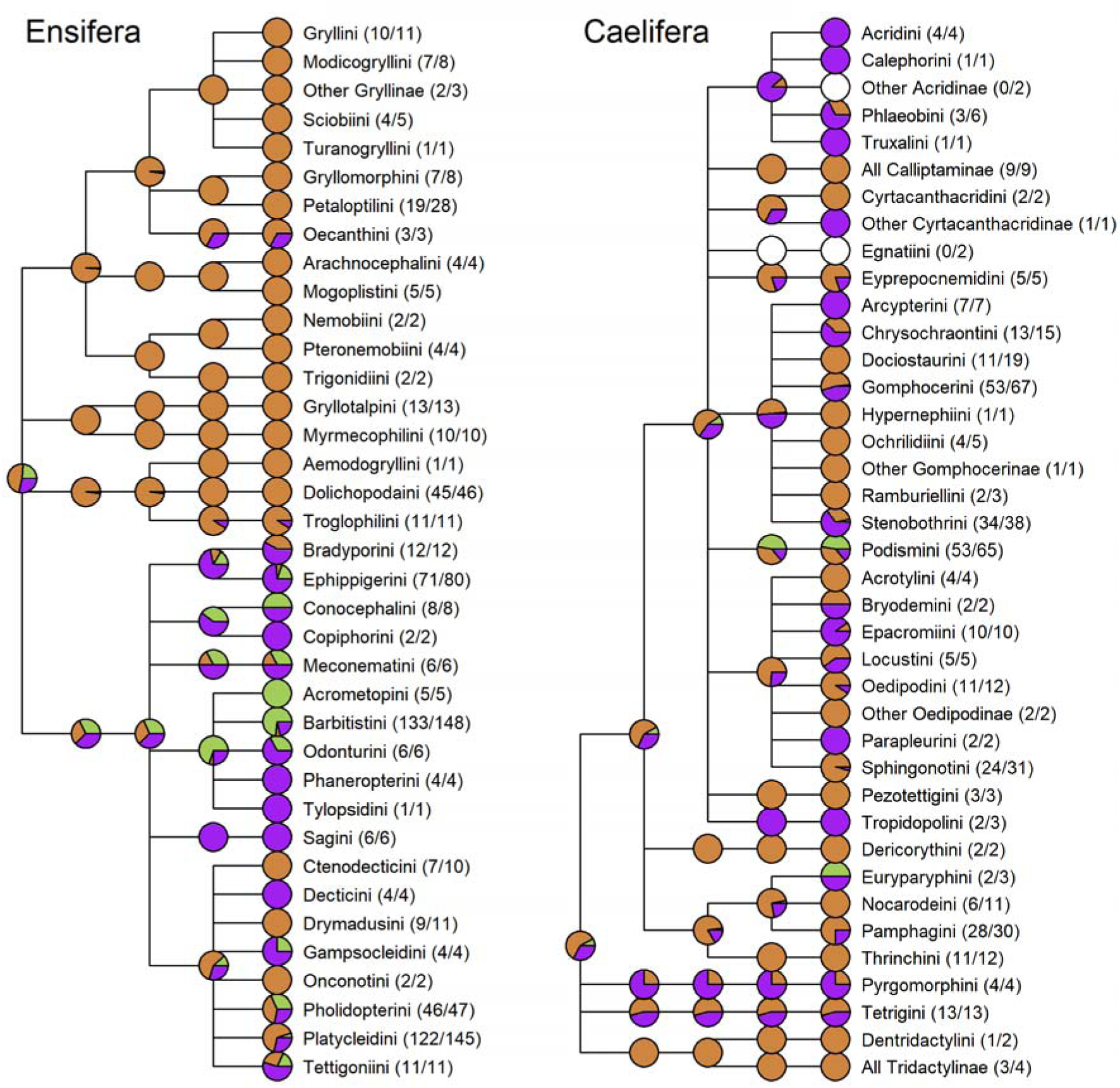
Taxonomic distribution of monomorphic brown (brown), monomorphic green (green) and polymorphic species (purple). Pie charts at each node represent the proportion of species within each clade that represent one of the states. Numbers (a/b) behind tribe names show (a) the number of species with morph information and (b) the total number of species per tribe. White pie charts represent clades with missing data for all species.

Polymorphic species occur more frequently in clades that comprise monomorphic green species than in those with otherwise only monomorphic brown species, a pattern that held at the level of genera and tribes (Figure 3a). Clades (at the level of genera and tribes) that contain monomorphic brown as well as monomorphic green species, almost always also contain species that are green-brown polymorphic (with the exception of a single genus, Figure 3a). This pattern was significant when analyzed at the level of genera (χ^2^ _2_ = 17.5, P = 0.0001) and at the level of tribes (χ ^2^_2_ = 9.6, P = 0.0081).

**Figure 3:**
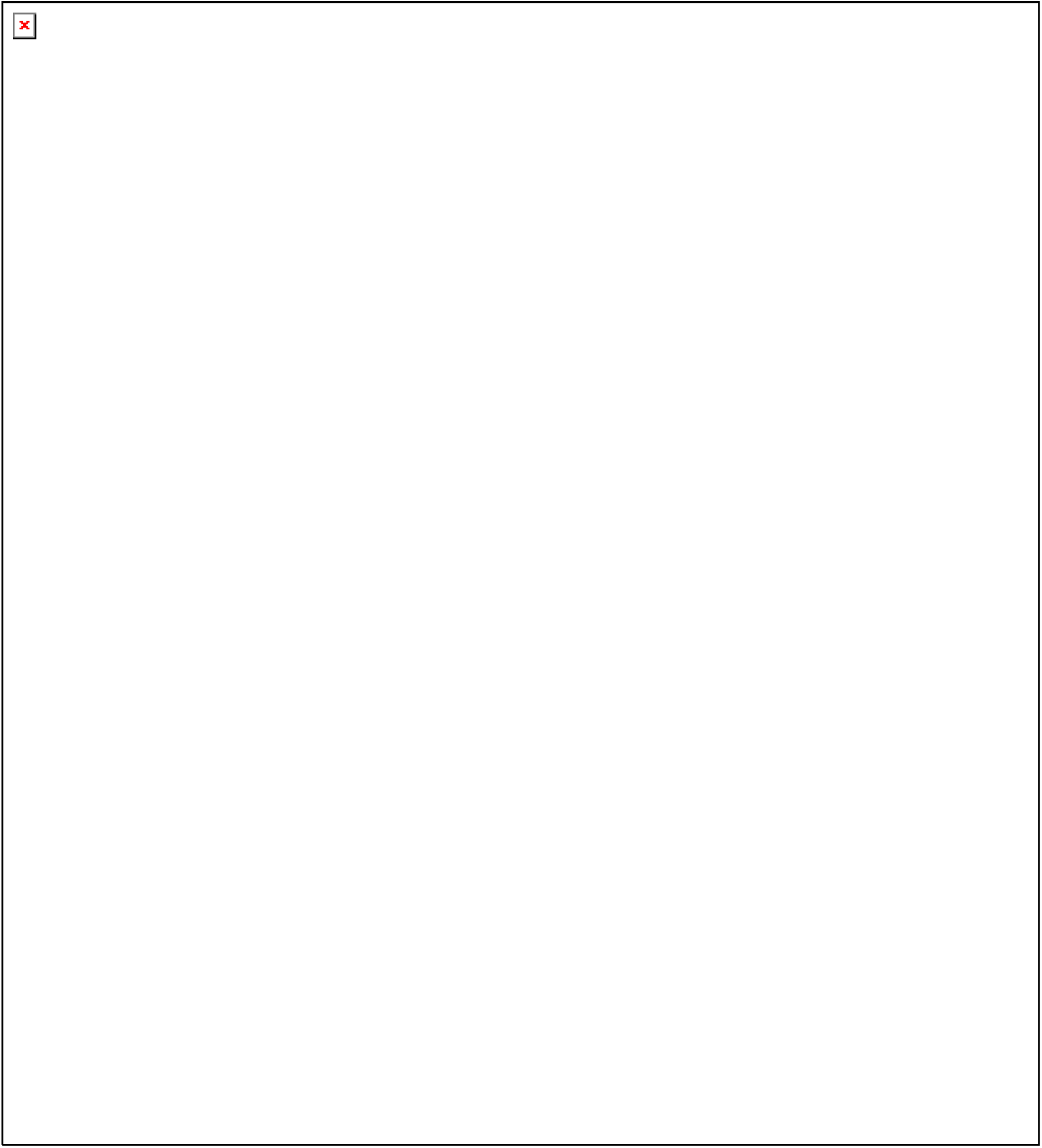
(a) Representation of polymorphic species in clades of species that are otherwise either entirely green, entirely brown, or with a mix of brown and green species. Filled bars show that number of clades with more than one species, while shaded bar sections include the number of clades with a single species. (b) Relative abundance of green and brown morphs across polymorphic species. Truncated bars on the sides represent monomorphic species. Note that the ratio of monomorphic brown to monomorphic green species is about 2.7:1.

### Relative abundances of color morphs in polymorphic species

Among color-polymorphic species, the less abundant morph is still common in 32%, regular in 36%, rare in 25%, and exceptional in 6% of the cases (Figure 3b). The relative abundance of the two morphs varies among color-polymorphic species and is skewed towards cases where green morphs is more common than the brown morph (Figure 3b). Particularly instructive are those cases in which the rare morph occurs at low frequencies. In 56 out of 73 cases where one of the morphs is rare this is the brown morph (χ ^2^ _1_ = 20.8, P < 10 ^-5^). In 10 out of 18 cases where one morph occurs exceptionally, this is the brown morph in otherwise green species (χ ^2^ _1_ = 0.22, P = 0.64). While the ratio 10:8 is almost equal, the ratio of purely brown to purely green species (thus the pool of species in which exceptional morphs can arise) is highly skewed towards brown (ratio 2.8:1).

### Geographic distribution

The occurrence of brown, green, and polymorphic species follows a geographic pattern. While north and central Europe host about 80% polymorphic species, polymorphic species compass only 40% of the species in southern Europe (Figure 4). The geographic distribution is not simply caused by the dominance of particular genera, since numbers are similar when looking at the proportion of genera and tribes that include polymorphic species. Both monomorphic green and, in particular, brown species are proportionally more abundant in southern Europe.

**Figure 4:**
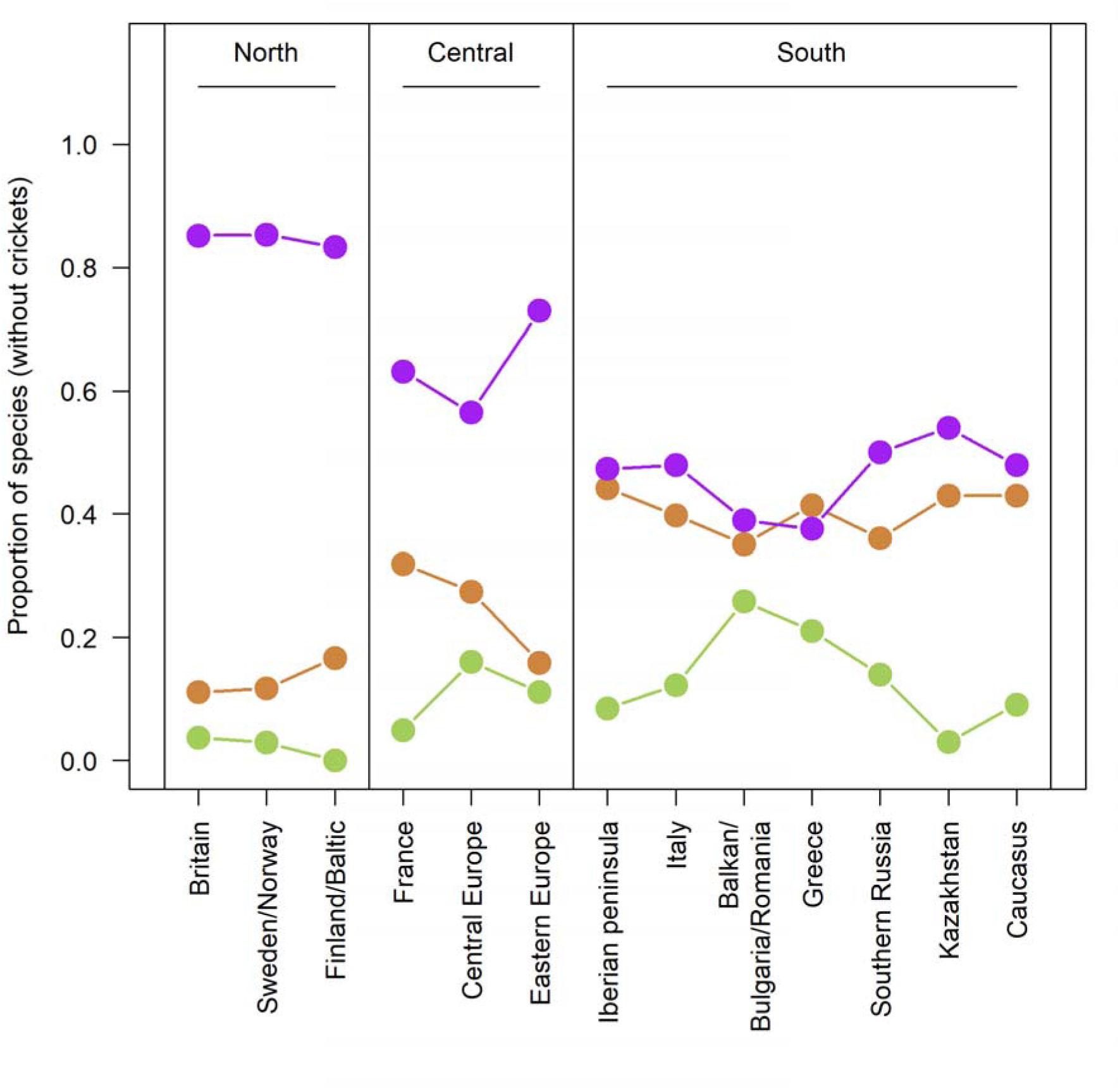
Geographic distribution of monomorphic brown (brown), monomorphic green (green) and polymorphic species (purple). The plot shows the proportion of species of each type in each region with more widely distributed species contributing to multiple categories. Crickets have been excluded, because they are very species rich in the south and are (almost) exclusively brown, so they would also strengthen the pattern that is evident in the graph.

### Distribution across habitats

Highest proportions of polymorphic species (about 60%) occur in marshes and moist grasslands (Figure 5a). Within grasslands, the proportion of polymorphic species declines from moist to dry grasslands, while the proportion of brown morphs increases. Alpine grass- and shrubland also host relatively high proportions of polymorphic species (45%, Figure 5a). Bushlands, forest margins and woodlands host a high proportion of green species and intermediate numbers of polymorphic species. Open and rocky habitats are dominated by brown morphs with relatively low numbers of polymorphic and hardly any monomorphic green species. Patterns are similar when looking at higher levels of taxonomic aggregation.

**Figure 5:**
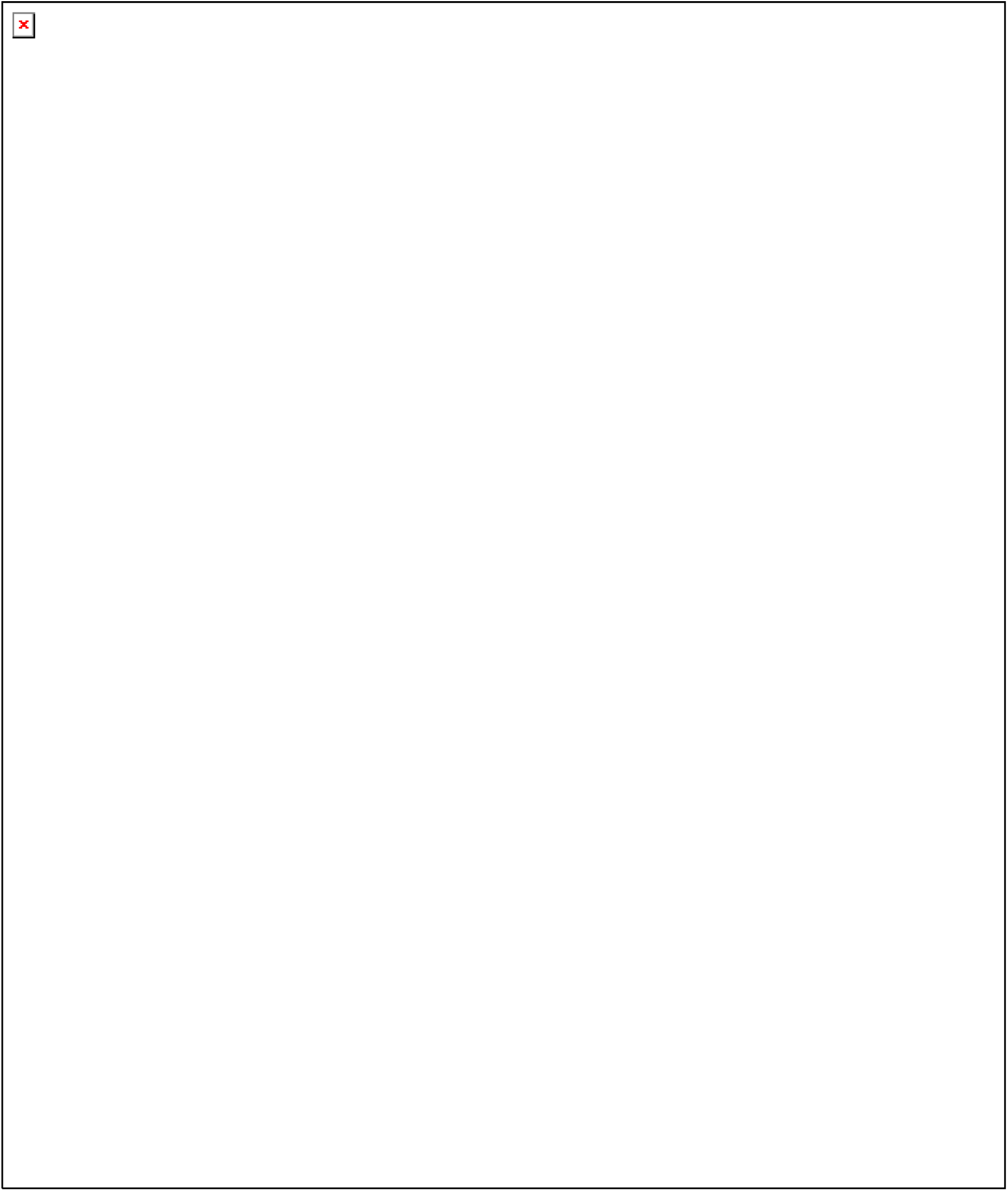
(a) Habitat distribution of monomorphic brown (brown), monomorphic green (green) and polymorphic species (purple). The plot shows the proportion of species of each type in each habitat. (b) Habitat use, diurnality and seasonal activity of monomorphic brown (brown), monomorphic green (green) and polymorphic species (purple). The plot shows the proportion of species of each type in each category.

Highest proportions of polymorphic species occur among species that dwell in bushes and trees or in higher vegetation (Figure 5b). The proportion of polymorphic species is also high among those species that live in lower vegetation and that regularly visit open ground. Quite to the contrary, predominately ground-dwelling species are usually monomorphic brown with low proportions of polymorphic species. Subterranean species are almost exclusively monomorphic brown. The proportion of polymorphic species is highest among diurnal species, while species that are active day and night and even more species that are exclusively nocturnal show lower proportions of polymorphic species (Figure 5b). Polymorphic species occur mostly during spring and summer, while species that overwinter are predominantly brown (Figure 5b).

### Species diversity, range sizes, and habitat diversity

There is no relationship between the proportion of polymorphic species and species richness within genera (Figure 6a,b) or at other taxonomic levels. However, there is a marked increase in the proportion of polymorphic species with increasing geographic range size (quantified as the number of geographic areas in which a species occurred) (Figure 6c). Similarly, there is a clear trend for increased polymorphism in species that occur in multiple habitat categories and can thus be considered more ecologically widespread (Figure 6d).

**Figure 6:**
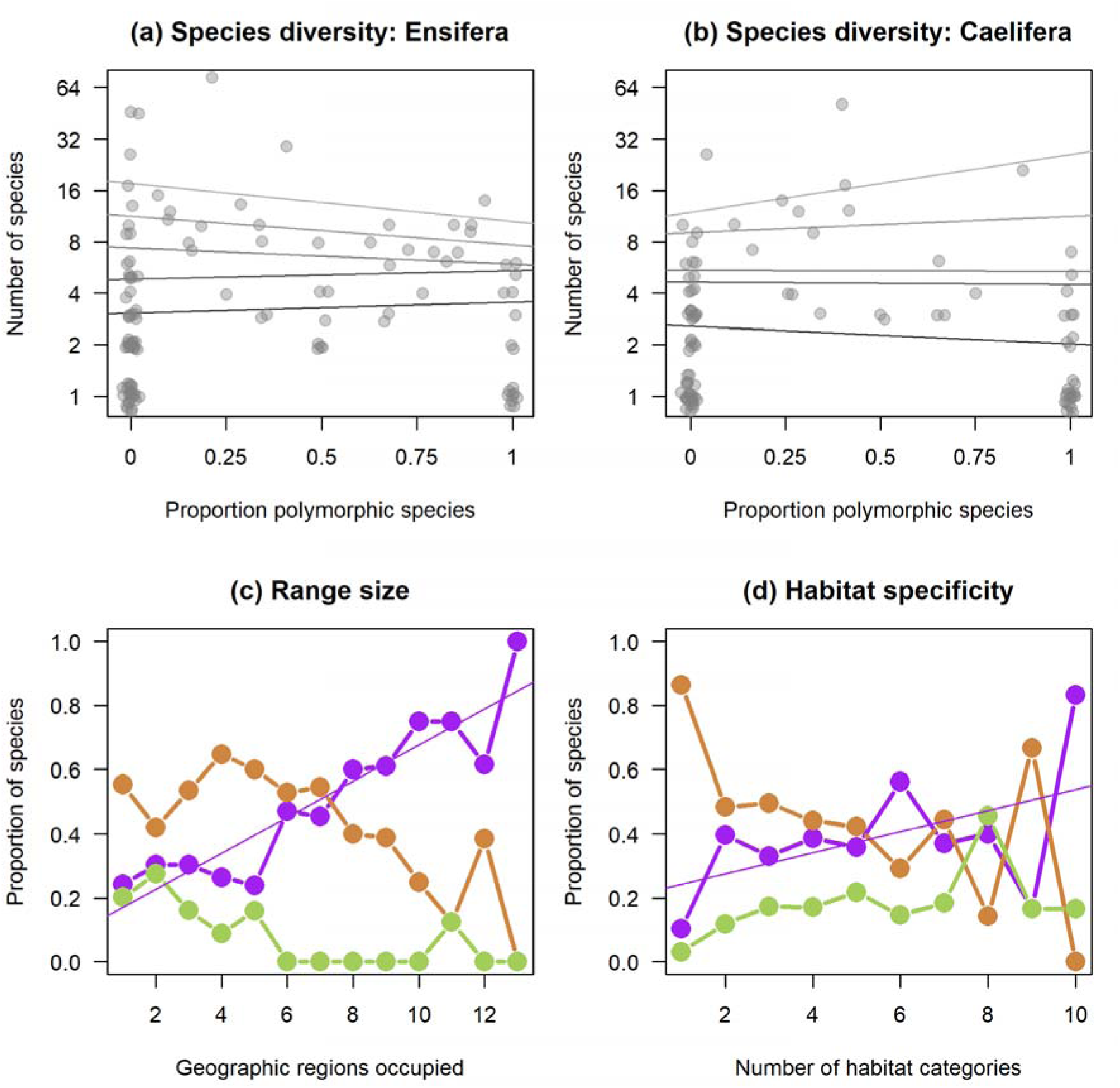
Correlations of species diversity, geographic range size and range of occupied habitats with intra-specific color morph diversity. (a, b) Number of species per genus and proportion of polymorphic species within (sub)genera. Each dot represents one (sub)genus. Points are partly overlaid, slightly jittered along both axes and darker dots show the overlay of multiple genera. Regression lines refer to increasing filtering from (dark to light) all species, minimum 2, 4, and 8 species. None of the regression slopes is significantly different from zero (P > 0.20). (c) Proportion of monomorphic brown (brown), monomorphic green (green) and polymorphic species (purple) in relation to the number of geographic regions inhabited by a species. The regression line for the proportion of polymorphic is statistically different from zero (P < 10^−5^). (d) Proportion of monomorphic brown (brown), monomorphic green (green) and polymorphic species (purple) in relation to the number of habitat categories inhabited by a species. The regression line for the proportion of polymorphic is not statistically different from zero (P = 0.14).

## Discussion

The data illustrate the high prevalence of the green-brown polymorphism in the European orthopteran fauna. This polymorphism is widespread geographically, phylogenetically, and across most habitats. The ratio of monomorphic brown, polymorphic green-brown, and monomorphic green species was about 5:3:2. The bias towards brown species and the dominance of polymorphic over green species might suggest that brown variants tend to be selected for in most habitats. However, the asymmetry is also consistent with a biased loss of green pigmentation in that the ability to produce green is more frequently lost than gained. The data also show evidence that within clades, it is more common to find polymorphic species when there are also green species than in clades that contain only brown species. This is consistent with the hypothesis, that the ability to produce the green pigment is limiting and more easily lost than gained.

Color polymorphisms have been implicated in favoring speciation (Gray and McKinnon 2007; West-Eberhard 1986). The European Orthoptera data does not show an association of taxonomic diversity with intraspecific polymorphisms. This could be a matter of phylogenetic resolution and the lack of a well-resolved and dated phylogeny. However, the current evidence is in no way suggestive of increased speciation in polymorphic orthopterans. Interestingly, polymorphic species tend to be more widespread, both geographically and also ecologically in terms of a broader range of habitats occupied. Causation in these cases can go either way, with either polymorphism favoring successful colonization or heterogeneous selection in widespread species favoring polymorphisms.

### A role of biased mutation

Several lines of evidence suggest that the green-brown polymorphism is sourced from loss-of-function mutations of a functional green pigmentation pathway. Frist, the frequency distribution of color morphs within species is skewed towards green individuals being more common in polymorphic species. In particular, cases of rare and exceptional morphs are more common in predominantly green populations. Examples of this are represented by species of the genera *Tettigonia, Phaneroptera* and *Conocephalus*, that are typically bright green, but show pale yellowish individuals at low frequencies. Second, polymorphic species are relatively more common in groups that contain green species than in clades that contain only brown species. Also, polymorphic species are almost universally present in groups that contain green as well as brown species. Third, breeding data suggest a dominant inheritance of the green variant at least in gomphocerine grasshoppers (Gill 1981b; Schielzeth and Dieker 2020; Winter et al. 2021). Some species are phenotypically plastic and can facultatively switch between the presence and absence of green colors (see discussion below). However, while this adds another level of complexity, it does not rule out that loss-of-function mutations play an important role. In particular, it is not clear if all individuals of a plastic species are able to switch color. Phenotypically plastic species could thus consist of a mix of families that are capable of producing green pigments, while others are limited to the expression of brown color. Only the former are expected to produce color-switching individuals. An obvious, yet untested, prediction of the biased loss-of-function hypothesis is that in polymorphic species there are more non-plastic families that are brown than non-plastic families that are green.

Based on these overall patterns, I suggest that a biased loss of green pigmentation with simultaneous selection for green variants in vegetated habitats creates polymorphic states and this creates a form of mutation-selection balance across species, even though the polymorphisms – once arisen – might well be maintained by balancing selection within species. Loss-of-function mutations play an important role in evolution (Behe 2010) and have been shown to contribute to the corolla color polymorphism in flowering plants (Whittall et al. 2006). This seems to be different from melanism in animals, where different non-synonymous substitutions in a few major genes seem to produce melanistic morphs (Hoekstra 2006; Majerus and Mundy 2003; Manceau et al. 2010). Molecular opportunities for loss-of-function mutations are manifold, thus can arise independently and repeatedly in different groups. The genetic basis of the color polymorphism is thus predicted to be different across different groups of orthopterans.

The pigment that produces the green color in orthopterans is supposedly biliverdin (Fuzeau-Braesch 1972) and in *Locusta migratoria* it has recently been shown that β-carotene plays a role in color transitions between green and black (Yang et al. 2019). However, it is not known if this applies to all species and biochemical analysis would help elucidating if the pigments are similar across all species. In most groups, the difference between green and brown morphs is in the presence or absence of green pigments. However, in some groups, the pigmentational basis could be different. In some species of *Barbitistes* or *Poecilimon*, for example, green colors appear to be covered by black coloration to a variable degree and it is not clear if green pigments are truly absent in very dark individuals. It is only those species in which the green-brown polymorphism might be considered continuous, while in the vast majority of species it is clearly discrete.

### Possible selective agents

The geographic distribution and the distribution of the polymorphism across habitats suggests a role for environmental filtering or natural selection in shaping morph ratios. Green morphs (both in a monomorphic state and in the form of polymorphism) are strongly underrepresented in dry, bare and underground habitats. For open ground species in dry habitats this is apparently related to crypsis, since green morphs would be more conspicuous in open, largely non-green habitats. For cave-dwelling, subterraneous and nocturnal species it need not be selection itself that is favoring brown variants, but green pigmentation could simply be lost because it is no longer selected for. Clades that have lost green pigmentation might even find themselves in an evolutionary trap in which a preference for a hidden lifestyle is favored because of loss of crypsis in most vegetated areas.

One of the main candidate mechanisms that can favor green variants is crypsis in vegetated habitats. Most orthopterans feed on leaves of grasses, herbs or shrubs (Ingrisch and Köhler 1998) and are exposed to predation from visual predators, such as birds, lizards and predatory (or parasitic) insects (Ingrisch and Köhler 1998; Ramme 1951). Visual predators thus represent potential selective agents, and selection for crypsis on vegetated patches is a potential promotor of the green morphs. This will favor the persistence of green pigmentation even if the green pigment is frequently lost by loss-of-function mutations.

The widespread distribution of the green-brown polymorphism calls for a rather general explanation, likely including some in the form of balancing selection. However, the selective agents that maintain coloration in a polymorphic state within populations are less easy to identify. It seems unlikely that recent admixture creates transient polymorphic states or that a large number of species is in transition from one state to another. There are only few cases of geographic variation in morph ratios that I am aware of (e.g. in *Chorthippus biguttulus* or *Decticus verrucivorus* green morphs increase with latitude, but populations are polymorphic across the range). I am not aware of any cases in which the geographic distribution of the polymorphism would suggest a role of admixture of different populations such as from different glacial refugia.

In some species, there are geographic clines in morph ratios (Köhler et al. 2017) and these can create polymorphic states in migration-selection balance. Color mediated thermoregulation with thermoregulatory benefits in brown morphs (Köhler and Schielzeth 2020) could favor brown variants in some patches and, if this trades-off with crypsis, green variants in others. While this process might be relevant in some species, it is doubtful how this can serve as a general explanation across more than 250 polymorphic species. A plausible candidate that could impose negative-frequency dependent selection is predation by predators that develop search images for the most abundant prey (Bond 2007; Dukas and Kamil 2001). However, there are no known species-specific predators, such that search image developments in predators (such as birds or lizards) would act on the community levels of orthopterans. No community-level predation study has yet been conducted.

The association of green and polymorphic species with habitat and geographic range clearly shows that color is related to environment, suggesting that color may be under environment-specific selection. Yet, aside from habitats such as caves, bare ground and rocks, the distribution of the frequency of color polymorphisms is also consistent with cross-species mutation-selection balance fueling the polymorphism. It remains an open question why the green-brown polymorphism seems to be prone to loss-of-function mutations and how many such variants exist across and within species. Once a polymorphism arises in green populations, the polymorphism can be maintained by balancing selection around some equilibrium that differs with the ecology of a species. However, with high mutation rates, morph ratios within species could also follow a random walk, drifting between mutation pressure producing brown and directional selection favoring green morphs. The relative role of directional and balancing selection will almost certainly differ between species.

### Other variants

While my main analysis is focused on the discrete green-brown polymorphism, grasshoppers are also remarkable for other color variation. Several open-habitat species vary between pale-grey, almost blackish, reddish and yellowish (Baños-Villalba et al. 2018; Edelaar et al. 2017; Ergene 1952; Peralta-Rincon et al. 2017). Bright purple or pink variants occur regularly in a number of species (Rowell 1972), but are never dominant in any species of the European fauna. Almost all species are variable in darkness. Most of this variation is based on variable amounts and ratios of different melanins (Fuzeau-Braesch 1972), but the variation appears continuous in most cases. However, there are pattern polymorphisms in a number of species, such as among Tettrigidae, in which these polymorphisms have a genetic basis (Ahnesjö and Forsman 2003; Fisher 1930; Karlsson et al. 2009). Lesser known pattern polymorphisms include pied variants (Dieker et al. 2018) among many species of Gomphocerinae (about 41% in my survey) and Oedipodinae (32%) and in one species of Pamphaginae (3%). Pied morphs seem to be absent from Ensifera. Many species, in particular of Gomphocerinae, occur in a dorsal striped variant that also affects the distribution of green coloration (Dearn 1990; Köhler 2006; Köhler et al. 2017; Richards and Waloff 1954). However, striped morphs come in very different variants and it is difficult to quantify the homologous striped variants.

## Conclusions

Overall, the results show the phylogenetically, geographically and ecologically widespread occurrence of the green-brown polymorphism in European orthopterans. With a representation in 30% out of more than 1,000 species of the European fauna, this represents one of the most penetrant phenotypic polymorphs in any group of organisms. Ecological and geographic patterns suggest an influence of selection and/or environmental filtering on the occurrence of polymorphisms. Interestingly, however, the patterns also suggest that biased loss-of-function mutations contributed to the green-brown polymorphism and that polymorphic states are primarily sourced from green rather than brown states.

